# Text Mining Drug-Protein Interactions using an Ensemble of BERT, Sentence BERT and T5 models

**DOI:** 10.1101/2021.10.26.465944

**Authors:** Xin Sui, Wanjing Wang, Jinfeng Zhang

## Abstract

In this work, we trained an ensemble model for predicting drug-protein interactions within a sentence based on only its semantics. Our ensembled model was built using three separate models: 1) a classification model using a fine-tuned BERT model; 2) a fine-tuned sentence BERT model that embeds every sentence into a vector; and 3) another classification model using a fine-tuned T5 model. In all models, we further improved performance using data augmentation. For model 2, we predicted the label of a sentence using k-nearest neighbors with its embedded vector. We also explored ways to ensemble these 3 models: a) we used the majority vote method to ensemble these 3 models; and b) based on the HDBSCAN clustering algorithm, we trained another ensemble model using features from all the models to make decisions. Our best model achieved an F-1 score of 0.753 on the BioCreative VII Track 1 test dataset.

## I. Introduction

Drug-protein interactions play an important role in biology. A great volume of articles on this topic are published every year, but only a few are manually annotated. It is thus important to develop an automated annotation system for these interactions in biomedical text.

The previous BioCreative challenge hosted a similar chemical-protein interaction track (1) where participants were asked to classify these interactions into various categories. The best system developed by (2) was able to achieve an F-1 score of 0.641. Recent advancements in text mining, especially the emergence of transformer based models, have improved the result significantly. The authors of Pubmed BERT (3) for example, reported an F-1 score of 0.772 in their work.

## II. Methods

### A. Data Processing

#### 1) The DrugProt Corpus for BioCreative VII Track 1

In the DrugProt Corpus (4), domain experts manually annotated the followings in the titles and abstracts of selected PubMed articles: i) all chemical and gene mentions; and ii) all relations between them corresponding to a set of 13 relation types. Their annotation criteria are available online.

The corpus is split into 3 subsets: training, development, and test. The test set contains 750 golden standard records where entities and relations are annotated manually, as well as another 10,000 background records with automatic annotations. The introduction of such huge background records, as the organizers wrote on their website, is for the following 3 purposes: i) to prevent participants from manually correcting predictions; ii) to generate a silver standard predicted relations; and iii) to test the capabilities of participating models on handling larger datasets. The organizers also released a large-scale test data with millions of abstracts and entities to further test the scalability of participating models. Table 1 shows the statistics of the above 4 datasets.

**TABLE I.**
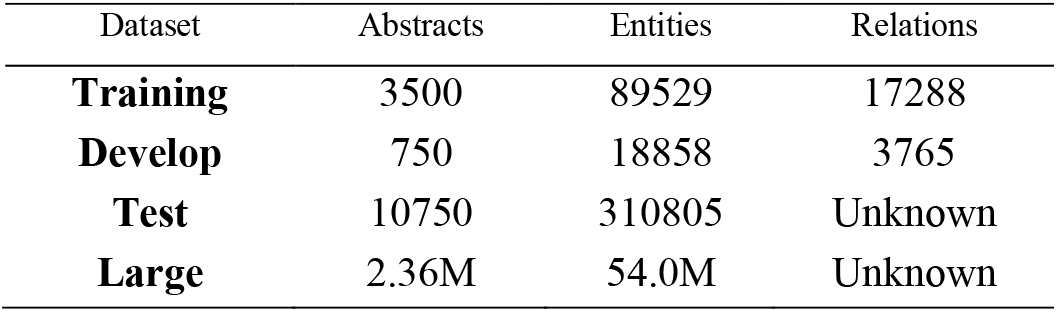
DrugProt Dataset Statistics

| Dataset | Abstracts | Entities | Relations |
| --- | --- | --- | --- |
| **Training** | 3500 | 89529 | 17288 |
| **Develop** | 750 | 18858 | 3765 |
| **Test** | 10750 | 310805 | Unknown |
| **Large** | 2.36M | 54.0M | Unknown |

In this article, we use (PMID, chem_arg, gene_arg) to reference the interaction between a chemical with label chem_arg and a gene with label gene_arg, in the PubMed article with PMID. We call such triplet an instance. When showing the text of an instance in this paper, we use *italic* and wrap the chemical and gene mentions of our interest with ** and $$ respectively. For example: *\*\*AG1478** inhibited EGF-induced $$MMP-9$$ expression*.

We discovered several interesting patterns in the dataset. First, while most annotated relation types are unique, there are some cases where a pair of gene and chemical entities in an article is annotated with multiple relation types. For example, (23064031, T6, T26): *\*\*NFD** suppressed EGF-mediated protein levels of c-Jun and c-Fos, and reduced $$MMP-9$$ expression and activity, concomitantly with a marked inhibition on cell migration and invasion without obvious cellular cytotoxicity*. The instance above is labeled with both INDIRECT-DOWNREGULATOR and INHIBITOR. Of the 17288 true relations in the training set, 430 have multiple labels. We discovered further that 28 of the 430 duplicates have the same interaction type. These are basically the same entry recorded twice in the dataset. Due to the small size of such occurrences, we do not consider the classification task as multi-label and proceed with training classifiers that predict only 1 class for each instance.

Second, we found 5697 overlapping entities in the training dataset. We say a pair of entities is overlapping when an entity is part of the other entity in a sentence. Note that his definition excludes entity pairs that have partial overlapping. We found only 5 instances that have a true interaction. In the 7th negative rule concerning CEMs (chemical Entity Mentions) that are part of a GPRO (Gene and Protein Related Object) mention in the official ChemProt annotation guideline (1), the organizers stated that these relations should not be annotated unless “there is an independent, separate mention of the same CEM in the same sentence, where it is explicitly mentioned that this CEM acts as a substrate of the GPRO.” Based on our observation of the data and this annotation rule, we ignore chemical-protein pairs that are overlapping when we process the data.

#### 2) Processing Pipeline

Our data processing pipeline consists of the following steps.

##### a) Step 1: Sentence tokenization

Sentence tokenization is the process of splitting a paragraph, an abstract in our application, into sentences. We use the NLTK tokenizer (5) fine-tuned with extra abbreviations and entities that should not be split. The extra info was first gathered by our previous team (6) in the BioCreative VI challenge and augmented this time by analyzing the tokenization results on the training dataset.

##### b) Step 2: Making false cases

While true interactions were annotated in the provided dataset, false ones were not. To generate false cases for each sentence, we first collected all the gene and chemical mentions. Then for each combination of gene and chemical we assign a “NOT” label to it if its relation is not annotated in the dataset. The underlying assumption is that the relation annotation is complete: all the interacting pairs were already annotated in the dataset.

##### c) Step 3: Entity name masking

Since we were training a classifier that predicts relation type based on the semantics of the sentences, we masked the chemical and gene names of all instances with special tokens: chem_name and gene_name respectively. This is to prevent the model from taking a short path and learn their relation type based on the names instead of the sentence semantics. We also tried masking the names of other named entities in the sentences but found the difference of model performance was negligible.

##### d) Step 4 Splitting dataset

During our internal testing, we split the provided training set into training and validation sets using a 4:1 split with 80% of the articles in the training set and the remaining 20% in the validation. Note that such split is based on articles, not instances, to prevent information leak. Were we to split the dataset based on instances, some instances that are from the same sentence would be split into different subsets, which would leak information and reduce the model’s generalizability when predicting the labels for new articles.

After testing internally using our own splits, we re-trained our models to submit results using the following 2 splits: i) using a 4:1 split on the training set as training and validation set; and ii) using a 4:1 split on the combination of training and development set as training and validation set.

##### e) Step 5 Data Augmentation

After training the BERT model, we obtained a subset of training data that the trained model predicted wrong. For every sentence, we used StanfordNLP to get the shortest path between chem_name and gene_name, and then randomly deleted one word not in the path. We labeled them the same as the originals, and augmented training datasets with these sentences in all our models.

### B. Models

In this work we used 3 models: i) BERT, ii) sentence BERT, and iii) T5. All these models are based on transformers. In this section, we briefly introduce the architectures, pre-trained models we used, post-processing steps if any, and the optimal hyper-parameters we used after fine-tuning the models.

#### 1) BERT model

The BERT model (7) is a transformer encoder model that stacks multiple layers of transformers on top of one another. We refer the readers to the original BERT paper (7) for details. For each sentence, we took the output of the classification token [CLS] from the last layer and passed it through a linear layer and a *tanh* activation function for the pooler output. We then applied a dropout layer and another linear layer to obtain the logits of each class. Finally, we used a sigmoid function to obtain the class probabilities.

Since the introduction of the BERT model, there have been multiple pre-trained BERT models in the biomedical domain, including BioBERT (8) and PubMedBERT (3). BioBERT is a continual pretraining from the original BERT and they share the same vocabulary. PubMedBERT however, pretrains everything from scratch including the vocabulary and thus performs better in downstream tasks for PubMed documents. We also observed during our experiments that PubMedBERT performed better than BioBERT. Specifically, the PubMedBERT version that is pre-trained on both the abstracts and full-text articles performed the best.

The BERT model was fine-tuned for 10 epochs with early stopping using the cross-entropy loss and the AdamW optimizer with a learning rate of 6e^−6^. These parameters were tuned on the validation set to optimize the F-1 score.

#### 2) Sentence BERT model

The sentence BERT model (7) we used has the same model architecture as the BERT model for generating token embeddings. After using the average over all token embeddings (mean pooling) to calculate sentence embedding, we transformed it by using a fully connected layer with 256 output dimensions and a *tanh* activation function to generate our final sentence embedding.

We used the same pre-trained weights as the BERT model as the initial values, and pre-trained our model for 10 epochs with early stopping to fine-tune the model. The model was fine-tuned with the AdamW optimizer with a 2e^−5^ learning rate with the batch hard triplet loss.

After fine-tuning the sentence embeddings, we used the HDBScAN (9) algorithm to cluster training data and used the k-nearest neighbor for classification.

#### 3) T5 model

The T5 model (10), short for Text-to-text Transfer Transformer, is a transformer encoder-decoder model that outputs text for every text input. It can deal with various tasks in one model, such as sentence translation, question answering, and classification. In our application, we considered all relations as *texts* and fine-tune the model to predict them for all sentences. We refer readers to the original paper for the model architecture, pre-training objectives, and model performance on downstream tasks.

We used SciFive-Large (11), a T5 model pre-trained on large biomedical corpora as our initial weights. We then trained the model for 10 epochs with early stopping, using the cross entropy loss and the AdaFactor optimizer with a learning rate of 0.001. The AdaFactor optimizer is known to be memory efficient, and it enables us to train the large model on a cluster of 2 GPUs with 12 GB memory each after splitting the model evenly.

### C. Ensemble Methods

In addition to training all the models individually, we also experimented with various ideas of training ensembles of them.

The first and simplest idea is using the majority vote of the three models. When all the three models predict differently, we use the result from the one that yields the highest F-1 score in the training data.

We also experimented with the idea of training ensemble models based on the clustering results from the Sentence BERT model and HDBSCAN. If the performances of the models vary from cluster to cluster, such ensemble methods can learn this pattern and adjust the weights of the models dynamically according to the cluster of any given sentence. Inspired by this idea, we implemented two ensemble models: i) simple ensemble, and ii) trained ensemble.

For the simple ensemble method, we assigned each cluster a model, which yielded the best accuracy over all the sentences in the cluster for the training data. We found using the accuracy metric instead of the F1 score to select models yielded the better result in our experiments. When making predictions, we first predicted the cluster of the sentence and then used the model assigned to that cluster to make the final prediction.

For the trained ensemble method, we extracted the following features from our models (because the T5 model was added in the last week of the competition, we did not have enough time to incorporate its results into the ensemble method): i) last layer features of the BERT model; ii) class probabilities of the BERT model; iii) cluster ID from the HDBSCAN; and iv) class label from k-NN classification. We trained an ensemble model of i) XGBoost (11); ii) Logistic regression; iii) Extra Trees classifier; and iv) Random Forest classifier, to predict one among the following four scenarios: i) both BERT and Sentence BERT predicted wrong; ii) Only BERT predicted the label right; iii) Only Sentence BERT predicted the label right; and iv) both models predicted the label right. During inference, for sentences that fall into the first scenario, we used the predicted label from the model that yielded the better F-1 score overall.

Figure 1 shows the diagram of our submitted models.

**Fig. 1.**
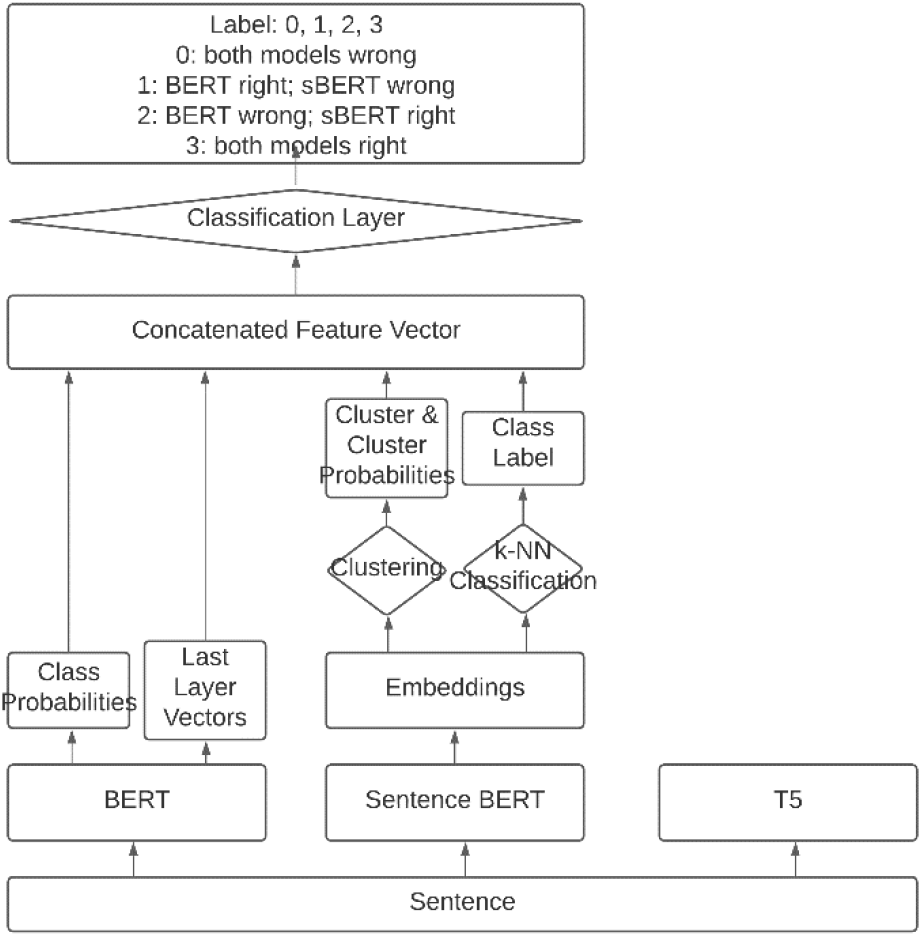
Diagram of Submitted Models

### D. Software

We implemented the BERT, sentence BERT and T5 models using transformers (12), sbert (13), and both both transformers and TensorFlow (14) respectively, and ensemble models using scikit-learn (15). Our code is available at github.com/luckynozomi/ChemProt-Biocreative

## III. Results

### A. Main Track

Besides the first run we submitted predictions from the model trained with 80% of training set and validated with the remaining 20% (the first train-validation split), all the others are trained with 80% of the combined training and validation sets and validated with the remaining 20% (the second train-validation split). We also trained a Sentence BERT model with the second split but did not include it in the submission because we could only submit 5 results. We also did not submit the result from the simple averaging ensemble because it was inferior to the majority vote method on the validation set.

The five runs we submitted are: 1. BERT model with the first train-validation split; 2. BERT model with the second train-validation split; 3. T5 model with the same train-validation split as Run 2; 4. Majority Vote of the BERT, Sentence BERT and T5 models; and 5. Trained ensemble of the BERT and Sentence BERT models. The results are in Table 2. Table 3 lists the detailed granular results by relation type.

**TABLE II.** Performance Metrics of Our Submitted Models Against the Average

| Submission | Precision | Recall | F1-validation | F1-test |
| --- | --- | --- | --- | --- |
| **1. Bert Split 1** | 0.702 | 0.786 | 0.721 | 0.742 |
| **2. Bert Split 2** | 0.659 | 0.824 | 0.747 | 0.732 |
| **3. T5** | 0.719 | 0.760 | 0.748 | 0.739 |
| **4. Majority Vote** | 0.754 | 0.751 | 0.764 | 0.753 |
| **5. Trained Ensemble** | 0.633 | 0.633 | 0.753 | 0.633 |
| **All Participants (mean ± std)** | 0.643±0.196 | 0.629±0.247 | | 0.620±0.232 |

**TABLE III.** Detailed Granular Results by Relation Type of the Majority Vote Ensemble

| Relation-Type | Precision | Recall | F1score |
| --- | --- | --- | --- |
| **ACTIVATOR** | 0.808642 | 0.784431 | 0.796353 |
| **AGONIST** | 0.755319 | 0.70297 | 0.728205 |
| **AGONIST-INHIBITOR** | 0 | 0 | 0 |
| **ANTAGONIST** | 0.86875 | 0.908497 | 0.888179 |
| **DIRECT-REGULATOR** | 0.767908 | 0.624709 | 0.688946 |
| **INDIRECT-DOWNREGULATOR** | 0.724551 | 0.796053 | 0.758621 |
| **INDIRECT-UPREGULATOR** | 0.711744 | 0.722022 | 0.716846 |
| **INHIBITOR** | 0.848309 | 0.835395 | 0.841802 |
| **PART-OF** | 0.573826 | 0.75 | 0.65019 |
| **PRODUCT-OF** | 0.591346 | 0.679558 | 0.632391 |
| **SUBSTRATE** | 0.679389 | 0.637232 | 0.657635 |
| **SUBSTRATE_PRODUCT-OF** | 0 | 0 | 0 |
| **AGONIST-ACTIVATOR** | 0 | 0 | 0 |

### B. Large Scale

Shown in Table 1, the large-scale dataset contains millions of abstracts and entities to predict. We started running predictions on September 23, 5 days before the submission deadline. We finished running the Sentence BERT model and submitted the result using 2 GPUs, while the BERT and T5 models required an extra day and did not finish.

Adapting our system to the large-scale dataset was straightforward. All we did was splitting the whole dataset into 100 slices, then run the whole pipeline on each of them. With 2 GPUs available, we let each run 50 slices. After obtaining prediction results on all 100 slices, we aggregated them into one single file and submitted it.

Our Sentence BERT model achieved a precision, recall and F-1 score of 0.71, 0.73 and 0.72, respectively.

## IV. Discussions

### A. Trained Ensemble Result

We believe there is a bug when submitting the trained ensemble result. In our validation set, it achieved a F-1 score of 0.753. Despite being inferior to the majority vote ensemble which had an F-1 score of 0.764, an F-1 score of 0.633 on the test dataset seems unlikely. However, due to the limitation of time, we have not located the problem yet.

### B. Analysis by Relation Types

Table 4 shows the confusion matrix of our best model, the majority vote model, on the validation set. The (i,j)th entry shows the number of entries with true label i and predicted as j. Due to the limitation of space, we only show the first 3 letters of each label. The bottom three rows are the precision, recall and F-1 score for each class.

**TABLE IV.** Confusion Matrix of the Majority Vote Ensemble on Validation Dataset

|  | ACT | AGO | AGO-ACT | AGO-INH | ANT | DIR-REG | IND-DOW | IND-UPR | INH | PAR-OF | PRO-OF | SUB | SUB_<br>PRO | NOT |
| --- | --- | --- | --- | --- | --- | --- | --- | --- | --- | --- | --- | --- | --- | --- |
| **ACT** | 220 | 0 | 0 | 0 | 0 | 2 | 0 | 18 | 6 | 0 | 0 | 0 | 0 | 67 |
| **AGO** | 1 | 91 | 0 | 0 | 0 | 8 | 0 | 0 | 6 | 0 | 0 | 0 | 0 | 16 |
| **AGO-ACT** | 0 | 3 | 0 | 0 | 0 | 0 | 0 | 0 | 0 | 0 | 0 | 0 | 0 | 0 |
| **AGO-INH** | 0 | 0 | 0 | 0 | 0 | 0 | 0 | 0 | 0 | 0 | 0 | 0 | 0 | 0 |
| **ANT** | 1 | 0 | 0 | 0 | 126 | 2 | 0 | 0 | 0 | 0 | 0 | 0 | 0 | 24 |
| **DIR-REG** | 2 | 0 | 0 | 0 | 0 | 344 | 1 | 0 | 20 | 7 | 1 | 5 | 0 | 102 |
| **IND-DOW** | 0 | 0 | 0 | 0 | 0 | 0 | 228 | 3 | 25 | 0 | 0 | 1 | 0 | 31 |
| **IND-UPR** | 7 | 0 | 0 | 0 | 0 | 0 | 11 | 181 | 5 | 0 | 0 | 1 | 0 | 63 |
| **INH** | 2 | 0 | 0 | 0 | 4 | 3 | 20 | 2 | 1240 | 0 | 0 | 4 | 0 | 141 |
| **PAR-OF** | 0 | 0 | 0 | 0 | 0 | 1 | 0 | 0 | 0 | 121 | 1 | 1 | 0 | 27 |
| **PRO-OF** | 0 | 0 | 0 | 0 | 0 | 0 | 0 | 0 | 0 | 0 | 145 | 12 | 0 | 29 |
| **SUB** | 0 | 0 | 0 | 0 | 0 | 2 | 0 | 0 | 5 | 1 | 5 | 317 | 0 | 72 |
| **SUB_PRO** | 0 | 0 | 0 | 0 | 0 | 0 | 0 | 0 | 0 | 0 | 1 | 5 | 0 | 0 |
| **NOT** | 71 | 14 | 0 | 0 | 17 | 155 | 71 | 46 | 183 | 89 | 70 | 157 | 0 | 8392 |
| **F** | 0.71 | 0.79 | 0.00 | 0.00 | 0.84 | 0.69 | 0.74 | 0.70 | 0.85 | 0.66 | 0.71 | 0.70 | 0.00 | 0.92 |

Class AGONIST-ACTIVATOR, AGONIST-INHIBITOR and SUBSTRATE_PRODUCT-OF have 0 F-1 scores because the numbers of samples in the dataset are very small. In fact, only 67 of 17288 instances have one of these labels. The lack of such instances makes it hard for the models to predict such cases. In fact, none of the instances in our validation set was predicted as any of these labels.

From the confusion matrix, we observed that there are not major confounding pairs among the true labels (all labels except NOT). Most of prediction errors occur between a true label and the NOT label. The most significant class of such mispredictions is PART-OF. We also observed this pattern in our clustering result. Of all the thirteen clusters, when we predicted the labels of all the validation data, PART-OF only appeared in one cluster. In that cluster, the only extra label is NOT. We infer that the Sentence BERT model, our sentence embedding algorithm, failed to separate these two classes.

We examined instances with PART-OF labels and none of the 10 nearest neighbors have the right prediction. We found several cases that are mislabeled. i) (18439678, T2, T20): *Both porcine $$TLR7$$ and TLR8 proteins were expressed in cell lines and were **N**-glycosylated*. ii) (10702256, T2, T11): *Removal of **N-** and O-linked oligosaccharides reduces the M(r) to approximately 160,000, suggesting that approximately 60% of the mass of SPACRCAN is $$carbohydrate$$*.

## References

1. Krallinger, M., Rabal, O., Akhondi, S.A., et al. (2017) Overview of the BioCreative VI chemical-protein interaction Track. Proceedings of BioCreative VI workshop.

2. Peng, Y., Rios, A., Kavuluru, R., et al. (2018) Extracting chemical-protein relations with ensembles of SVM and deep learning models. Database, 2018.

3. Gu, Y., Tinn, R., Cheng, H., et al. (2020) Domain-Specific Language Model Pretraining for Biomedical Natural Language Processing. 1, 1–24.

4. Miranda, A., Mehryary, F., Luoma, J., et al. (2021) Overview of DrugProt BioCreative VII track: quality evaluation and large scale text mining of drug-gene/protein relations. Proceedings of the seventh BioCreative challenge evaluation workshop.

5. Wagner, W. (2010) Steven Bird, Ewan Klein and Edward Loper: Natural Language Processing with Python, Analyzing Text with the Natural Language Toolkit. Language Resources and Evaluation, 44.

6. Lung, P.Y., He, Z., Zhao, T., et al. (2019) Extracting chemical-protein interactions from literature using sentence structure analysis and feature engineering. Database, 2019.

7. Devlin, J., Chang, M.-W., Lee, K., et al. (2018) BERT: Pre-training of Deep Bidirectional Transformers for Language Understanding..

8. Lee, J., Yoon, W., Kim, S., et al. (2019) BioBERT: a pre-trained biomedical language representation model for biomedical text mining. 1–8.

9. McInnes, L., Healy, J. and Astels, S. (2017) hdbscan: Hierarchical density based clustering. The Journal of Open Source Software, 2.

10. Raffel, C., Shazeer, N., Roberts, A., et al. (2020) Exploring the limits of transfer learning with a unified text-to-text transformer. Journal of Machine Learning Research, 21.

11. Phan, L.N., Anibal, J.T., Tran, H., et al. (2021) SciFive: a text-to-text transformer model for biomedical literature. CoRR, abs/2106.0.

12. Wolf, T., Debut, L., Sanh, V., et al. (2020) Transformers: State-of-the-Art Natural Language Processing.

13. Reimers, N. and Gurevych, I. (2020) Sentence-BERT: Sentence embeddings using siamese BERT-networks. EMNLP-IJCNLP 2019 - 2019 Conference on Empirical Methods in Natural Language Processing and 9th International Joint Conference on Natural Language Processing, Proceedings of the Conference.

14. Abadi, M., Barham, P., Chen, J., et al. (2016) TensorFlow: A system for large-scale machine learning. Proceedings of the 12th USENIX Symposium on Operating Systems Design and Implementation, OSDI 2016.

15. Pedregosa, F., Varoquaux, G., Gramfort, A., et al. (2011) Scikit-learn: Machine Learning in Python. Journal of Machine Learning Research, 12, 2825–2830.

